# The Genome of the Human Pathogen *Candida albicans* is Shaped by Mutation and Cryptic Sexual Recombination

**DOI:** 10.1101/310201

**Authors:** Joshua M. Wang, Richard J. Bennett, Matthew Z. Anderson

**Affiliations:** Department of Molecular Microbiology and Immunology, Brown University, Providence, RI, 02912, USA; Department of Microbiology, The Ohio State University, Columbus, OH, 43210, USA; Department of Microbial Infection and Immunity, The Ohio State University, Columbus, OH, 43210, USA

## Abstract

The opportunistic fungal pathogen *Candida albicans* lacks a conventional sexual program and is thought to evolve, at least primarily, through the clonal acquisition of genetic changes. Here, we performed an analysis of heterozygous diploid genomes from 21 clinical isolates to determine the natural evolutionary processes acting on the *C. albicans* genome. Consistent with a model of inheritance by descent, most single nucleotide polymorphisms (SNPs) were shared between closely related strains. However, strain-specific SNPs and insertions/deletions (indels) were distributed non-randomly across the genome. For example, base substitution rates were higher in the immediate vicinity of indels, and heterozygous regions of the genome contained significantly more strain-specific polymorphisms than homozygous regions. Loss of heterozygosity (LOH) events also contributed substantially to genotypic variation, with most long-tract LOH events extending to the ends of the chromosomes suggestive of repair via break-induced replication. Importantly, some isolates contained highly mosaic genomes and failed to cluster closely with other isolates within their assigned clades. Mosaicism is consistent with strains having experienced inter-clade recombination during their evolutionary history and a detailed examination of nuclear and mitochondrial genomes revealed striking examples of recombination. Together, our analyses reveal that both (para)sexual recombination and mitotic mutational processes drive evolution of this important pathogen in nature. To further facilitate the study of genome differences we also introduce an online platform, SNPMap, to examine SNP patterns in sequenced *C. albicans* genomes.

**AUTHOR SUMMARY:** Mutations introduce variation into the genome upon which selection can act. Defining the nature of these changes is critical for determining species evolution, as well as for understanding the genetic changes driving important cellular processes such as carcinogenesis. The fungus *Candida albicans* is a heterozygous diploid species that is both a frequent commensal organism and a prevalent opportunistic pathogen. Prevailing theory is that *C. albicans* evolves primarily through the gradual build-up of mutations, and a pressing question is whether sexual or parasexual processes also operate within natural populations. Here, we determine the evolutionary patterns of genetic change that have accompanied species evolution in nature by examining genomic differences between clinical isolates. We establish that the *C. albicans* genome evolves by a combination of base-substitution mutations, insertions/deletion events, and both short-tract and long-tract loss of heterozygosity (LOH) events. These mutations are unevenly distributed across the genome, and reveal that non-coding regions and heterozygous regions are evolving more quickly than coding regions and homozygous regions, respectively. Furthermore, we provide evidence that genetic exchange has occurred between isolates, establishing that sexual or parasexual processes have transpired in *C. albicans* populations and contribute to the diversity of both nuclear and mitochondrial genomes.

## INTRODUCTION

A wide variety of genetic events contribute to the evolution of eukaryotic genomes. In asexual cells, haploid genomes evolve via the accumulation of point mutations as well as undergo recombination events that drive DNA expansions/contractions (indels). Heterozygous diploid genomes also have the capacity to experience loss of heterozygosity (LOH) events, in which genetic information is lost from one of the two chromosome homologs. In addition, both haploid and diploid genomes may experience large-scale chromosomal changes such as gross rearrangements, acquisition of supernumerary chromosomes or other forms of aneuploidy [1, 2].

Many eukaryotic species also generate genetic diversity via sexual reproduction. Here, recombination between individuals provides an efficient mechanism for producing diverse progeny. Sexual reproduction can therefore promote adaptation to new environments more rapidly than asexual propagation [3, 4]. However, this comes at a fitness cost due to the associated energetic requirements and the fact that only 50% of parental alleles are passed on to single progeny [5-7]. Sex can also be detrimental by breaking up beneficial allelic combinations [5, 8]. Facultative sexuality, the ability to alternate between sexual and asexual forms of reproduction, promotes a flexible lifestyle that can accelerate adaptation in response to environmental pressures [4, 9].

Sexual reproduction has been extensively studied in the *Saccharomyces* clade, where the model yeast *Saccharomyces cerevisiae* divides mitotically but can also undergo mating and meiosis to generate recombinant progeny. The related *Candida* clade includes some of the most important human fungal pathogens encountered in the clinic [10, 11], although the *Saccharomyces* and *Candida* clades diverged from one another ~235 million years ago [12]. The most clinically-relevant *Candida* species is *C. albicans* that, like all *Candida* species, was originally designated an obligate asexual organism. However, mating of diploid cells has been observed in the laboratory and produces tetraploid cells that return to the diploid state via a parasexual process of concerted chromosome loss (CCL) [13-16]. Mating requires that *C. albicans* cells undergo a phenotypic transition from the sterile “white” state to the mating-competent “opaque” state [17]. Conjugation of opaque cells can occur via heterothallic or homothallic mating [18], and recombination during CCL involves Spo11, a conserved ‘meiosis-specific’ factor involved in DNA double-strand break formation across diverse eukaryotes [14, 19].

Clinical isolates of *C. albicans* exhibit a largely clonal population structure despite the potential for recombination via parasexual reproduction [20, 21]. Multilocus sequence typing (MLST) separates *C. albicans* isolates into 17 clades although previously described incompatibility between MLST haplotypes and individual mutations suggests that recombination may act to generate new allelic variants [22]. Analysis of a limited number of haploid mitochondrial loci also reveals allelic mixtures that suggest recombination may have occurred within *C. albicans* populations [23, 24]. However, despite these observations, *C. albicans* is still commonly assumed to be an asexual species that does not undergo mating or recombination in nature [25, 26]. Prior studies focused on a subset of genomic loci and present conflicting evidence regarding the role of recombination in shaping *C. albicans* evolution [20, 22-24, 27], which can now be addressed by a detailed analysis of full genome sequences.

In this work, we examined evolutionary patterns in 21 sequenced *C. albicans* isolates that represent different clades, different sites of infection in the host, and different countries of origin [28, 29]. Our analyses provides a detailed picture of how mutational events drive evolution of the diploid *C. albicans* genome. We reveal that mutations preferentially accumulate in heterozygous regions of the genome, and that emergent SNPs and indels often cluster together. Moreover, we highlight isolates whose nuclear and mitochondrial genomes appear highly admixed and therefore display evidence of genetic contributions from multiple clades. These results establish that the *C. albicans* genome is a dynamic landscape shaped both by local mutations and large-scale rearrangements, and that sexual or parasexual mating has made a significant contribution to genotypic variation.

## RESULTS

The availability of whole genome sequencing data for 21 diverse *C. albicans* isolates [28] provided an opportunity to determine how genetic diversity is generated between strains in nature. The *C. albicans* diploid genome is ~14 megabases (Mb) and consists of eight chromosomes encoding ~6100 genes [28, 30, 31]. SNPs occur at a frequency of ~0.3% between chromosome homologs in the standard laboratory strain SC5314 (i.e., an average of 1 SNP every 330 bp) [28, 32]. Among the 21 isolates, we found SNP frequencies varied from 0.5% between closely related strains within Clade I to 1.1% between strains from different clades (Table S1). A previous phylogenetic reconstruction using 112,223 SNP positions found that most strains matched their previously assigned fingerprinting clades and MLST subtypes, with the exception of P94015 which clustered separately from other Clade I strains (Fig. 1A) [28]. Strong bootstrap values across the constructed phylogeny of these strains supports a primarily clonal lifestyle in which most polymorphisms are mutations consistent with inheritance by descent (Fig. 1B). Accordingly, SNPs and indels fit a nonrandom distribution across the 21 sequenced isolates χ^2^ (((SNPs; 20, N = 302641) = 83118, p < 2E-16, indels; 20, N = 19581) = 13825, p < 2E-16, Fig. S1).

**Figure 1.**
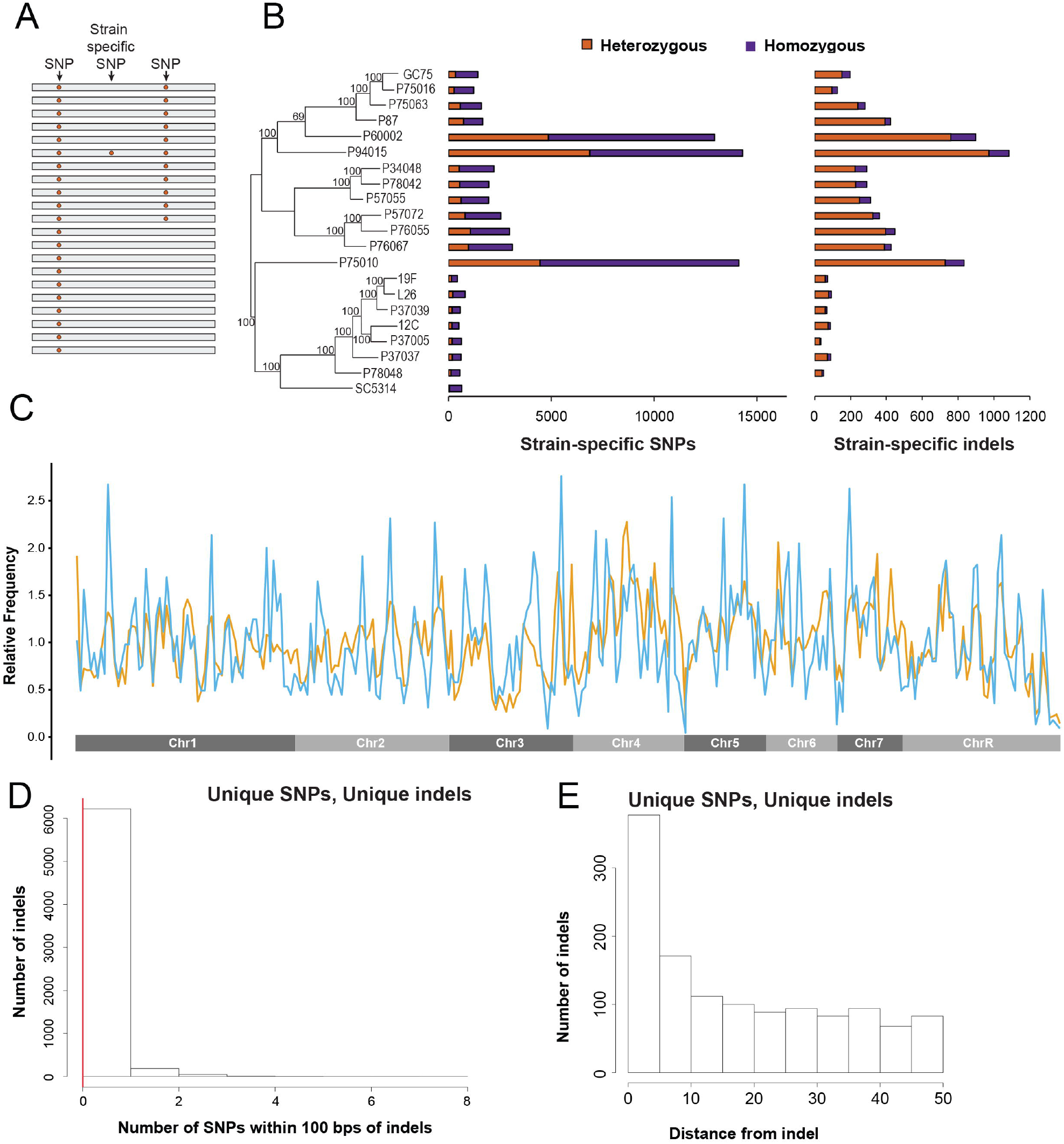
Distribution of polymorphisms among 21 clinical isolates of *C. albicans*. **A.** Number of heterozygous (purple) and homozygous (orange) strain-specific SNPs and insertion/deletions (indels) are plotted for each isolate. Clade designations for each isolate are color coded. **B.** There are two types of sequence variants among the set of 21 isolates; single nucleotide polymorphisms (SNPs) shared by multiple isolates and SNPs that are specific to individual strains. Variants encoded by multiple strains suggest origins in a common ancestor, whereas strain-specific polymorphisms likely arose specifically in individual strain backgrounds. **C.** Relative frequency of strain-specific SNPs (blue) and strain-specific indels (orange) were plotted across the genome using 5 kb sliding windows. **D.** Number of strain-specific SNPs within 100 bp of each strain-specific indel was plotted. The average number of strain-specific SNPs in an equal number of random 100 bp windows bootstrapped 1000 times is shown (red line). **E.** Distance to the nearest strain-specific SNP for each strain-specific indel was measured in a 100 bp window (with non-overlapping 10 bp intervals) surrounding the indels.

### Base substitutions in *C. albicans*

LOH events can distort the patterns of SNPs inherited from ancestral strains (Fig. S2). To help limit these confounding effects, we restricted most analyses to strain-specific SNPs and indels that are unique to individual strains. Approximately 25% of all SNP positions and 10% of all indel positions were strain-specific (66,086 and 6,474 events, respectively; Tables S2, S3). As expected, the number of strain-specific mutations increased with longer branch lengths from the nearest node in the phylogenetic tree (SNPs, *r_s_* = 0.60, p = 4.2E-3; indels, *r_s_* = 0.42, p = 0.055; Fig. 1B and Fig. S3). Correlation between these metrics of strain identity supports the use of strain-specific mutations in assessing mutational patterns.

In many eukaryotes, base-substitution mutations are biased towards transitions over transversions, although the cause of this bias is not completely clear [33]. In *C. albicans*, base substitutions also favored transitions over transversions for both strain-specific SNPs and total SNPs, χ^2^ ((11, N = 66086) = 18182, p < 2E-16 and (11, N = 302641) = 628000, p < 2E-16, respectively). The ratio of transitions to transversions was 2.21 for strain-specific SNPs and 2.50 for all SNPs (Fig. S4). Both coding and noncoding regions encoded more strain-specific transitions than transversions, although coding sequences were more biased than noncoding regions (2.74 versus 1.80, respectively). Base substitutions displayed a 1.39-fold bias towards introducing A/T instead of G/C for strain-specific SNPs that shrank to 1.03-fold when including all SNPs. The fact that substitutions favor transitions resulting in A/T suggests that this may contribute to the overall A/T richness of the *C. albicans* genome [31].

### Distribution of strain-specific polymorphisms across the *C. albicans* genome

Analysis of the global distribution of strain-specific SNPs revealed a bias against the accumulation of these mutations within protein-coding genes. Thus, most strain-specific polymorphisms (33,818 of 66,086 SNPs and 5,502 of 6,474 indels) were present within the 36.7% of the genome representing intergenic regions, suggesting that mutations in coding sequences are selected against (p = 9.71E-16; Fig. 2A,B). As a result, relatively few strain-specific SNPs were present within ORFs across the twenty-one sequenced strains (Fig. 2C). We found that 259 genes exhibited significantly greater SNP densities per nucleotide (nt) than the 0.004 SNPs/nt average for all *C. albicans* ORFs (Fig. 2C, Table S4). SNP densities within enriched genes were equal to or greater than the intergenic average (0.0066 vs. 0.0063, respectively). Protein-coding genes within this group lacked any enrichment for gene ontology (GO) annotations or pathways (Table S4). However, noncoding snoRNAs (small nucleolar RNAs) were significantly overrepresented among ‘faster-evolving genes’ by GO term analysis, χ^2^ ((2, N = 5) = 15.6, p = 7.90E-5; Fig. 2D). The five snoRNAs identified from GO enrichment had mutation rates greater than 0.02 SNPs/nt, significantly higher than that of the average rate of 0.0063 SNPs/nt within intergenic regions. Strain-specific polymorphisms clustered towards the 5’ end of the snoRNAs (Fig. 2E) and could contribute to variation in functional aspects of protein translation, although this possibility was not explored here.

**Figure 2.**
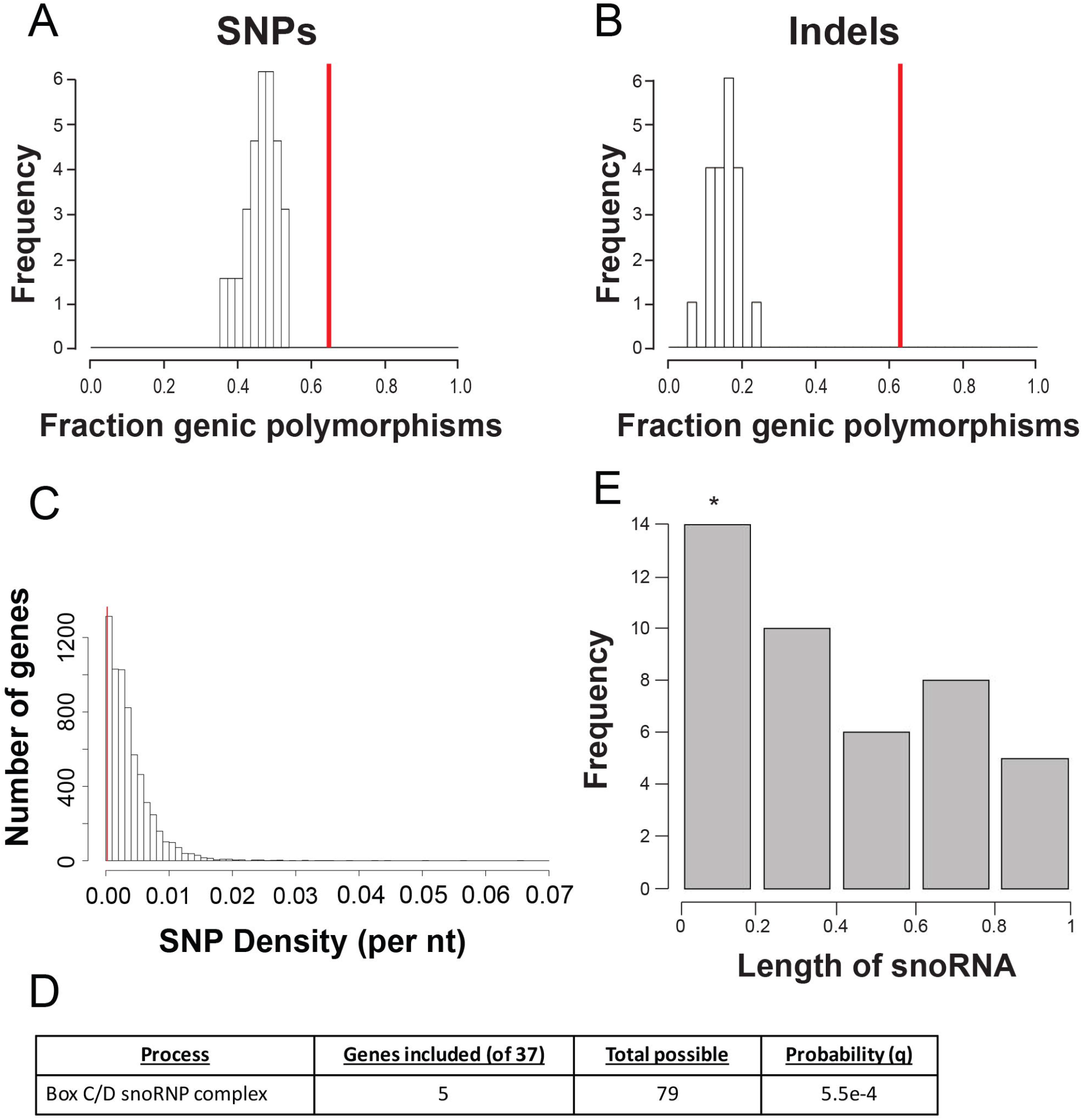
Strain-specific SNPs are enriched at intergenic positions and snoRNAs. The ratio of genic to intergenic strain-specific SNPs **(A)** and indels **(B)** was calculated for each of the sequenced isolates and falls below the fraction of the genome that encodes protein-coding genes (red line). **C.** The density of strain-specific SNPs from all sequenced isolates was measured for all genes in the *C. albicans* genome and plotted against the average SNP density for all isolates (red line). **D.** The placement of the SNPs was determined by breaking snoRNAs into five equal segments. SNPs were significantly enriched in the 5’ end of the RNAs. * denotes p < 0.01. **E.** Genes enriched for strain-specific SNPs were identified and functional enrichment was assessed using GO term analysis. snoRNAs were significantly enriched for strain-specific SNPs.

An inspection of strain-specific indels revealed that 3527 (54.5%) were deletions and 2948 (45.5%) were insertions. Indels ranged in size from 1 bp to 10 bp with the majority of longer events being insertions (Fig. S5). The frequency of both insertions and deletions decreased as mutations became larger, suggesting that smaller events occur more frequently or are less detrimental to the cell and therefore are retained more often. The incidence of ±3 nucleotide indels (21.9% of the total) was higher than that expected by chance. When indels were separated into genic or intergenic mutations, intergenic mutations followed a Poisson distribution centered on 0, whereas genic mutations were vastly overrepresented for ±3 nucleotide indels that do not shift the reading frame (Fig S5). Only ~15% of all indels fell within ORFs (p < 2.2E-16) suggesting that, as with SNPs, indels are selected against within coding sequences (Fig. 2B).

Indels have been commonly associated with specific genomic features such as repetitive sequences in other species [34, 35]. Across the sequenced *C. albicans* isolates, there was a total of 19,581 indel sites across the genome. Of these, 465 indel sites (2.37%) were located within annotated repetitive sequences (long terminal repeats (LTRs), major repeat sequences (MRSs), and retrotransposons). Total indels are therefore overrepresented within these repetitive features (two-tailed Brunner-Munzel (BM) test = 5.15, df = 182.05, p = 6.65E-7). Likewise, strain-specific indels were significantly enriched within repetitive features (47 of 6475; BM test = 13.004, df=182, p<2.2E-16). Both total and strain-specific SNPs also clustered within repetitive elements (BM test = 14.98, df = 315.62, p < 2.2E-16 and BM test = 12.26, df = 240.02, p < 2.2E-16, respectively). Thus, mutations within the *C. albicans* genome are enriched within repetitive regions similar to what has been observed in other species [34, 35].

Analysis of the genome-wide distribution of strain-specific SNPs and indels across the 21 genomes revealed that these mutation types showed significant clustering with one another (Fig. 1C) (Pearson, t = 11.64, df = 286, p < 2.2E-16). Multiple SNPs often occurred within 100 bp of an indel (Fig. 1D) as was confirmed via Sanger sequencing of selected regions (Fig. S6). Enrichment of SNPs was observed immediately adjacent to indels (within 10 bp) but not within indels (Wilcoxon test (W(1.79E7)), p < 2.2E-16; Fig. 1E). Three strains, P60002, P75010 and P94015, encoded a large proportion of strain-specific mutations reflective of their longer branch lengths in the phylogenetic tree, which could potentially skew the analysis (Fig. 1A). However, even after removing these three strains from the analysis and reducing the four major clades to three representative strains each, we still observed a significant association between SNPs and indels (Wilcoxon test (W(3.04E7)), p < 2.2E-16, Fig. S7A). This association highlights that distinct mutagenic events occur in close proximity to one another, and suggests that indel formation or the associated DNA repair processes may be mutagenic in *C. albicans*.

In some species, the introduction of indels can influence the observed mutational bias towards either transitions or transversions [35, 36]. To address this possibility in *C. albicans*, the transition:transversion ratio was determined for the ~500 strain-specific SNPs located within 10 bp of strain-specific indels. Although base substitutions still slightly favored transitions, the 1.17 transition:transversion ratio was significantly lower than the genome-wide average ratio of 2.21 (p = 5.87E-7). This is consistent with mutations close to indels exhibiting a reduced bias towards transitions over transversions due to recruitment of error-prone polymerases during DNA repair [35]. We therefore suggest that a similar mechanism operates in *C. albicans* and can account for the increased mutation rate adjacent to indels, as well as the local bias in the transition:transversion ratio.

### Association between LOH recombination events and base-substitution mutations

The previous study by Hirakawa *et al*. identified extensive loss of heterozygosity (LOH) tracts in the 21 sequenced *C. albicans* isolates [28]. Consequently, LOH breakpoints were mapped in each isolate and emphasis was placed on the distribution of LOH events around the mating type-like *(MTL)* locus on Chr5. The current study extends the analysis of LOH patterns in *C. albicans* genomes by determining if genome-wide patterns of LOH exist, and if there is an association between LOH tracts and other mutational classes such as base substitutions or indels.

LOH regions were defined in Hirakawa *et al*. using several parameters including contiguous 5 kilobase (kb) windows with a high frequency of homozygous SNPs (>0.4 events per kb; see Methods and [28]). Plotting the incidence of LOH for all chromosomes (Chr) in each of the isolates revealed a striking pattern, whereby the prevalence of LOH increased along each chromosome arm when progressing from centromere to telomere (Fig. 3A and Fig. S8). In fact, the overwhelming majority of all long-tract LOH regions (155 out of 170 regions larger than 50 kb) extended to the ends of the corresponding chromosomes (Table S5). This reveals that out of a total of 336 chromosome arms in the 21 isolates, 155 of these arms show evidence of having undergone a long-tract LOH event. LOH frequency decreased towards the centromeres and did not occur across centromeres except during LOH of whole chromosomes (Fig. 3B). Interestingly, LOH frequencies remained low across the entirety of the right arms of Chr2 and Chr4 (Fig. S8), suggesting that heterozygosity of loci on these arms may be maintained by selection. Aneuploidy did not significantly alter the frequency of heterozygous and homozygous intervals along aneuploid chromosomes relative to euploid chromosomes (p = 0.756).

**Figure 3.**
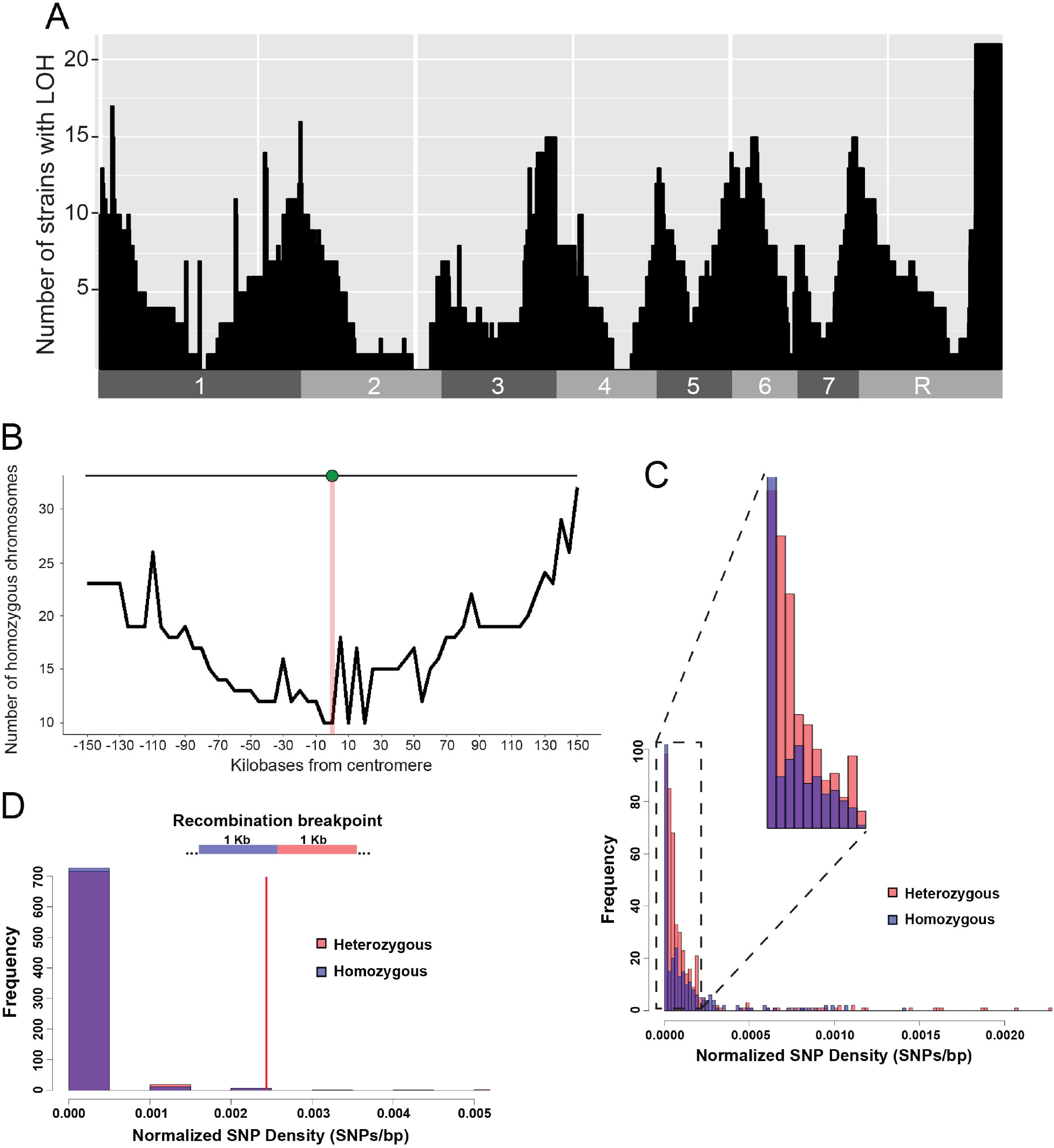
Loss of heterozygosity (LOH) events influence the *C. albicans* genome landscape. **A.** Number of strains that show LOH for 50 kb windows across the *C. albicans* genome aligned to their chromosomal position. **B.** Number of chromosomes that are homozygous for 5 kb windows within 150 kb of the centromere. **C.** SNP frequency for each of the 445 heterozygous (het) and 468 homozygous (hom) regions across the 21 sequenced genomes was plotted. Normalized SNP density is calculated using the number of SNPs within a het or hom region divided by the region length, and is significantly elevated for het regions (red) relative to hom regions (blue). **D.** One kb segments flanking the 745 LOH transition points were separated into their respective het and hom segments and assessed for SNP density. The genome average was calculated by randomly selecting an equal number of 2 kb windows across the genome and bootstrapping for 1000 iterations.

Several studies have revealed that mutation rates can be impacted by the underlying genomic context. For example, accumulation of SNPs was found to be increased in regions adjacent to indels in diverse eukaryotic species [36], and mutation rates were higher in heterozygotes than in homozygotes during meiosis [37]. We therefore examined mutational patterns in *C. albicans* genomes that are a mosaic of heterozygous and homozygous regions. We subdivided *C. albicans* genomes into heterozygous (het) or homozygous (hom) regions using defined criteria on all SNPs (see Methods), resulting in 468 het and 445 hom regions, respectively (Table S5). Het regions covered a total of 71.1% of the genome and hom regions 28.9%; het tracts were therefore considerably longer on average than hom tracts (~480,000 bp vs. 186,000 bp, respectively). Definition of het and hom regions using all SNPs allowed subsequent examination of the frequencies of <u>strain-specific</u> SNPs within these regions. Strain-specific SNPs comprise only 3% of all SNPs within these genomes and, therefore, do not contribute substantially to the designation of het and hom regions. Notably, het regions contained significantly higher frequencies of strain-specific SNPs than hom regions (1.4E-4 vs. 7.3E-5 SNPs/bp, respectively; BM test = −10.6, df = 786.6, p < 2E-16; Fig. 3C). Even after the exclusion of the three outlier strains, P60002, P75010 and P94015, and reducing the four major clades to three representative strains each, there were still significantly higher frequencies of strain-specific SNPs within het than within hom regions (BM test = −7.558, df = 377.0, p = 3.14E-13). Furthermore, all 21 isolates exhibited the same bias towards het regions containing more strain-specific SNPs than hom regions (two-tailed BM test = −1.11, df = 38.5, p = 0.28), indicating that mutations preferentially accumulate in het over hom regions during natural evolution of *C. albicans* isolates.

We note that heterozygous SNPs may have arisen in hom regions but, in some cases, been eliminated by a subsequent LOH event. As LOH can occur in one of two possible directions (due to loss of either homolog A or homolog B), we accounted for mutations potentially lost via LOH by doubling the number of homozygous, strain-specific SNPs within hom regions. Even with this adjustment, het regions still contained a greater density of strain-specific SNPs than hom regions (two-sided BM test = −8.74, df = 717.58, p<2.2E-16). The ~2-fold greater accumulation of polymorphisms in het over hom regions of the *C. albicans* genome shows parallels with the ~3.5-fold higher mutation rate observed in het vs. hom regions of the *Arabidopsis* genome during meiosis [37].

Sites close to recombination events, including LOH events, have been shown to be associated with elevated mutation rates in some species [36, 38-40]. To determine if there is an increased frequency of SNPs in regions proximal to LOH tracts in *C. albicans*, the density of SNPs at heterozygous-homozygous transition points was investigated. Analysis of the 745 identified transition regions included 1 kb of DNA on either side of the junction points between het and hom regions (with the latter inferred to represent LOH tracts). The SNP density within these transition regions was significantly lower than that in the rest of the *C. albicans* genome (one-sided BM test = −35.415, df = 748.8, p <2.2E-16; Fig. 3D). Furthermore, SNP density was similar on both het and hom sides of the LOH breakpoint. Thus, base substitutions appear to accumulate less frequently in regions proximal to het/hom breakpoints in the *C. albicans* genome. One caveat noted here is that this result may be influenced by difficulty in the identification of precise breakpoints between het and hom regions of the genome.

### Identity by descent during *C. albicans* evolution

A hallmark of phylogenetic reconstructions in asexual species is the ability to track the relatedness of isolates based on inherited polymorphisms [22, 41, 42]. Reconstruction often relies on a maximum parsimony model of ‘identity by descent’, in which more closely related strains share a greater percentage of shared polymorphisms (Fig. 4A). *C. albicans* SNP patterns generally follow identity by descent; these patterns matched the phylogenetic tree as evidenced by strong bootstrap support at almost all nodes [28], as well as a visual examination of SNP patterns within specific regions (Fig. 4B,C, and Fig. S9). Strikingly, however, certain regions of the genome exhibited clear violations of identity by descent (Fig. 4D,E, and Fig. S10). In heterozygous diploid genomes, these deviations could potentially arise through two mechanisms: (1) by sexual recombination between genetically distinct isolates, or (2) through multiple, independent LOH events that obfuscate the actual pattern of descent (Fig. S11). In the latter case, multiple LOH events could cause loss or retention of SNPs through homozygosis of one chromosome homolog or the other, thereby generating a subset of isolates that appear “recombinant”, i.e., appear to have intermixed genetic content from two different relatives. Such a history can sometimes be inferred by a comparison of SNP patterns within the region of interest in multiple extant strains (Fig. 4A).

**Figure 4.**
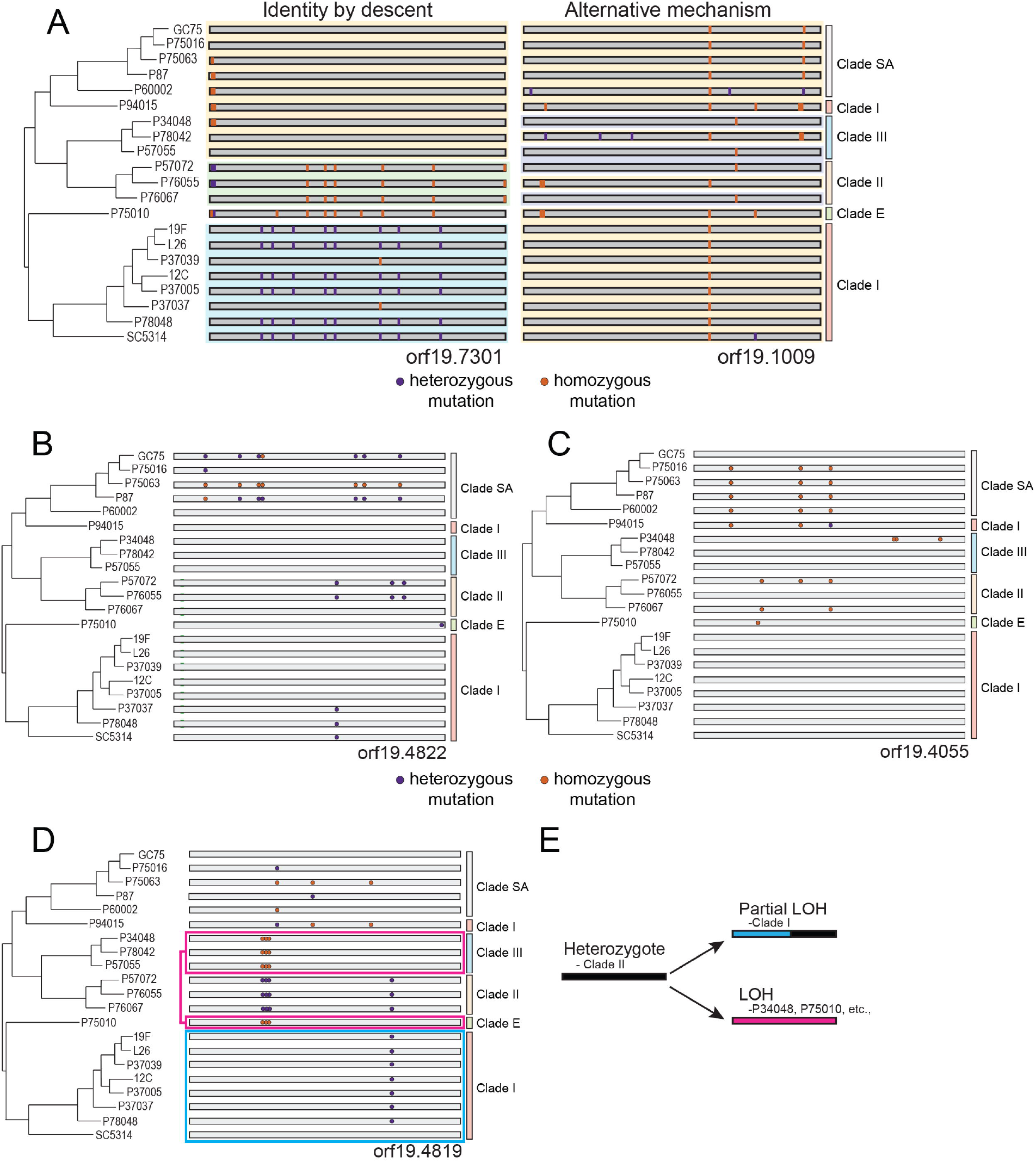
Mutational patterns following identity by descent and loss of heterozygosity are widespread. **A.** Two patterns of polymorphisms exist within sequenced genomes. One shows polymorphisms are phylogenetically congruent indicative of identity by descent, and the other shows polymorphisms that violate the phylogenetic relationship implicating mechanisms other than direct inheritance. Shading denotes similarity in overall SNP patterns. Heterozygous SNPs are purple and homozygous SNPs are orange. Polymorphisms for the set of *C. albicans* genomes are plotted for two loci (**B, C**) that display SNP patterns consistent with inheritance by descent. **D.** The polymorphism pattern is plotted for a locus that does not follow inheritance by descent. Similar genotypes are color-coded and connected to each other. **E.** A cartoon depicting LOH of heterozygotes in opposing directions provides the most parsimonious explanation for the observed SNP patterns.

We note that certain regions appear to have undergone divergent, short-tract LOH events across the 21 isolates, consistent with these events often occurring during asexual divisions (Fig. 4D,E, Fig. S10 and Supplementary Material). In line with this, we identified 514 non-overlapping 25 kb windows across the 21 sequenced isolates that do not encode similar polymorphisms to closely-related strains, and could represent regions that had experienced LOH. Interestingly, two strains, P60002 and P94015, contained 390 of these regions (75.9%), although only 214 of the 390 incongruent regions in these two strains (54.9%) overlapped with LOH tracts (hom regions) in these isolates. In contrast, the majority of the incongruent regions in all other strains (117 of 124 regions) overlapped with LOH tracts. This suggests that incongruence in polymorphisms in most strains likely results from divergent LOH events but that LOH does not obviously explain the majority of incongruent polymorphic patterns observed in P60002 and P94015.

### Evidence for recombination in natural isolates of *C. albicans*

Previous studies have provided conflicting messages regarding recombination in natural populations of *C. albicans* [20, 22-24, 27], and none have examined whole genome data for evidence of inter-clade mixing. We therefore examined the 21 sequenced genomes for mixed evolutionary histories. The similarity of genomic segments from each strain to the overall phylogenetic tree was compared by analysis of SNP patterns using 25 kb sliding windows. To aid visualization of SNP patterns we developed a custom interactive tool, SNPMap (http://snpmap.asc.ohio-state.edu/), which allows users to map the positions of individual mutations, mutation types, and het/hom tracts across user-defined regions of the 21 genomes.

Our analysis largely focused on two clinical isolates, P60002 and P94015, that have the weakest bootstrap support within the *C. albicans* phylogeny and that cluster with different strains when analyzed by MLST or DNA fingerprinting [28], suggesting their genomes may be recombinant. In support of this, examination of Chr4 in P94015 identified one region with clear homology to Clade I which was in close proximity to a region highly homologous to Clade SA (Fig. 5A, Table S6). The region with homology to Clade I (labeled P94015-A in Fig. 5A) shares a large number of SNPs that are present throughout Clade I but are absent in all other strains with the exception of P94015. Next to this region, a 1 kb segment (region P94015-B) lacks clear homology to any of the other sequenced isolates while, adjacent to this, the SNP pattern in P94015 is virtually identical to that of two Clade SA isolates (region P94015-C).

**Figure 5.**
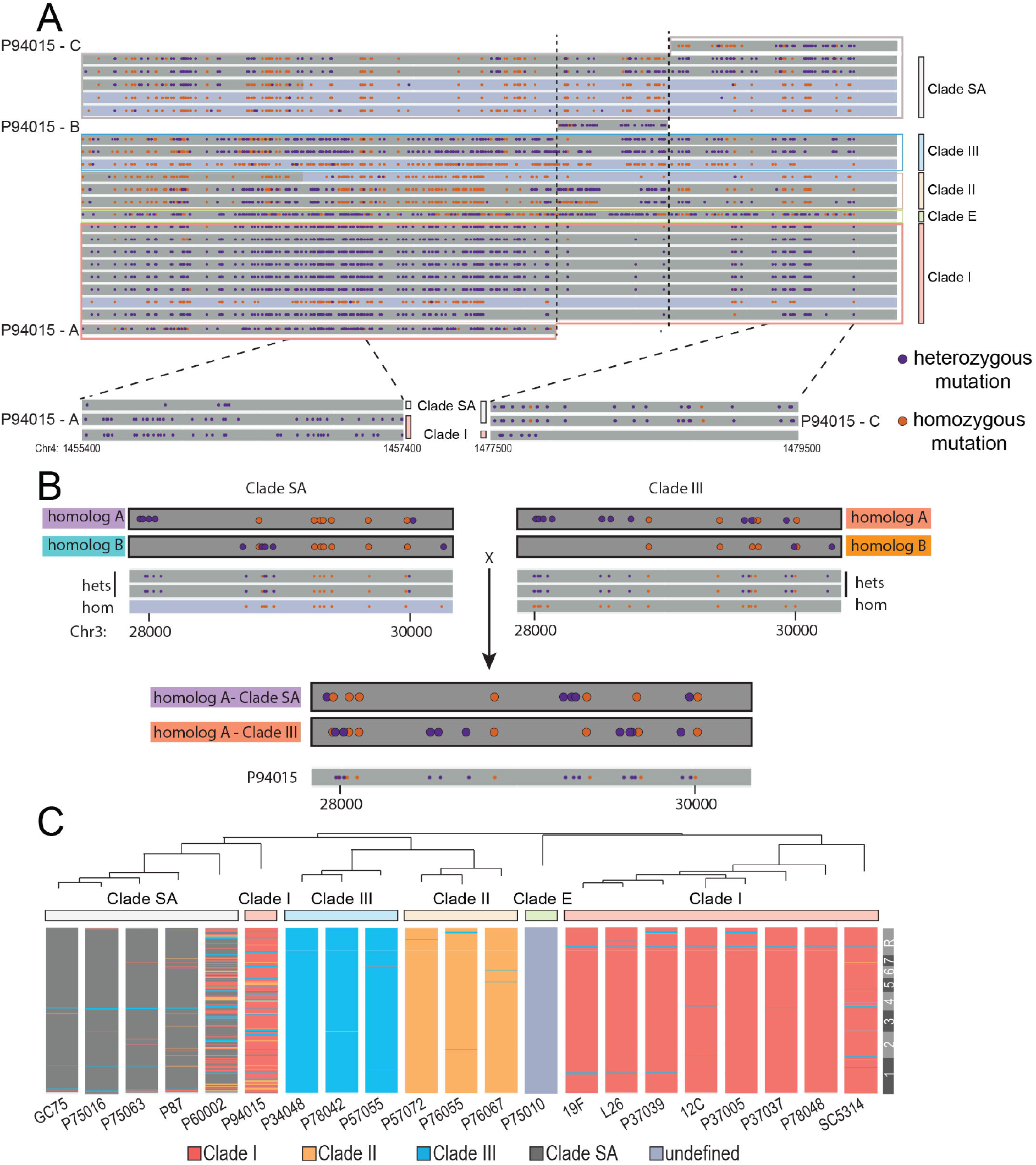
Evidence for recombination in *C. albicans* isolates. **A.** All SNPs are shown for a 20 kb region of Chr4 for the 21 sequenced genomes. For each strain, dark grey bars are heterozygous genomic regions while lighter bars indicate regions that are mostly homozygous. The SNP pattern in P94015 indicates one region with homology to Clade I (P94015-A) next to a region without clear homology to any specific clade (P94015-B) followed by a region with homology to Clade SA (P94015-C). **B.** SNPs for a 2.5 kb region of Chr3 were phased to individual homologs for Clade SA and Clade III by using LOH of a closely-related strain for reference. One homolog from each clade matched the exact SNP pattern in a ‘hybrid’ region present in P94015 (both the position of the SNP and the actual base substitution matched between P94015 and the Clade SA or Clade III homolog). **C.** Consensus SNP patterns for each clade were used to assess genome similarities between all isolates in 25 kb sliding windows. The closest match for each window was color-coded by clade. The SNP patterns for two strains, P60002 and P94015, contained regions assigned to multiple clades. dark grey-SA, blue-Clade III, mustard-Clade II, red-Clade I, light grey-no clade consensus.

Mating between isolates from different clades would be expected to generate hybrid DNA regions, with SNPs on one homolog of the recombinant strain matching those in one clade and SNPs on the other homolog matching those in a second clade. Identifying inherited SNPs following *C. albicans* mating in nature is complicated by the fact that, with the exception of SC5314 [32], haplotypes are not available for the 21 *C. albicans* genomes. Despite this, phasing of heterozygous SNPs for some isolates can be inferred using SNP patterns from related strain(s) that have undergone LOH for that region (Fig. 5B). The region that experienced LOH will only retain the SNPs that reside on the same homolog (i.e., those that are phased). Using this approach, we phased SNPs for chromosomal regions of closely related strains that are heterozygous in some isolates but homozygous in others. Multiple isolates that have undergone LOH for both alleles strengthen the confidence of phasing assignments within a given clade.

We applied this approach to a region on Chr2 in P94105 that contains polymorphisms identical both to those on homolog A of a Clade SA strain (12 of 12 SNPs are identical) and to those on homolog A from a Clade III isolate (15 of 15 SNPs are identical) (Fig. 5B). Both SNP positions and nucleotide identities are conserved across this hybrid region in P94015 when compared to the corresponding homologs from Clade SA and Clade III isolates. This region therefore provides a striking example of P94015 inheriting one homolog from a Clade SA strain and one homolog from a Clade III strain, and establishes a non-clonal origin for this isolate. Analysis of additional regions for isolates P94015 and P60002 provides support for the existence of multiple recombination tracts indicating that mixing has occurred between strains from different *C. albicans* clades (Fig. S12).

To examine global patterns of admixing among the set of 21 isolates, the distribution of all variant positions in each strain was compared to the consensus pattern for each clade using sliding 25 kb windows. The SNP patterns of most isolates resembled the consensus pattern for their assigned clade (98.5% of genomic windows matched their assigned clade), as expected for a population propagating clonally (Fig. 5C). In contrast, many regions within the P60002 and P94015 genomes showed homology to multiple different clades, producing highly mosaic genomes (Fig. 5C). Here, the genomes of P60002 and P94015 matched their assigned clades for only 58.3% and 76.7% of sliding windows, respectively (p = 1.14E-10). The majority of genomic regions in P94015 aligned with Clade I (genome is mostly red in Fig. 5C), whereas numerous segments aligned to regions from three other major clades (SA, II, and III). In the case of P60002, Clade SA regions made up the majority of the genome with a smaller number of regions matching Clade I or, to a lesser extent, Clade II. In line with this, the branchpoint leading to P60002 is the least well-supported node in the phylogenetic reconstruction of all 21 isolates [28]. The most parsimonious explanation for these highly mosaic genomes is that they are the products of mating and recombination between isolates from multiple *C. albicans* clades.

### Analysis of mitochondrial genomes in *C. albicans* isolates

Haploid mitochondrial genomes provide a more simplified context to search for evidence of recombination than heterozygous diploid genomes. In S. *cerevisiae*, mitochondrial genomes are biparentally inherited following mating, and recombination can occur between parental genomes prior to zygote division [43]. We therefore performed the first comparative analysis of global SNP patterns in *C. albicans* mitochondrial genomes using sequencing data from the set of 21 isolates. The mitochondrial genome in *C. albicans* is ~41 kb in size and a total of 1847 SNPs (and 0 indels) were annotated within the 21 isolates, with an average SNP density of 1 polymorphism every 476 bp. The mitochondrial genome was highly heterogeneous including areas of high SNP density (e.g., ChrM: 15000-20000) and regions devoid of polymorphisms (e.g., ChrM: 6000-12000) among sequenced isolates. Furthermore, of the 1847 annotated mitochondrial SNPs, only 39 were strain-specific in the set of 21 sequenced genomes (Table S2).

*C. albicans* mitochondrial genomes generally showed clade-specific SNP patterns that were again consistent with a clonal population structure, although resolution of SNP patterns was low due to relatively few clade-defining mitochondrial SNPs (Fig. 6). As with nuclear genomes, examination of mitochondrial genomes of P60002 and P94015 again showed clear evidence of inter-clade mixing. For example, the mitochondrial genome of P94015 contained regions that aligned with mitochondrial segments from both Clade SA and Clade II (Fig. 6A). Here, there are three polymorphisms that are Clade SA-specific on ChrM: 1-6000 (region P94015-A), and all three are present in P94015 (Fig. 6A). An additional two of the fifteen P94015 polymorphisms in this region are specific to this strain (Table S2). The remainder of the mitochondrial genome in P94015 (region P94015-B) encodes 144 polymorphisms matching the SNP pattern found in Clade II (with the exception of two strain-specific SNPs). Recombination between Clades I and II was even clearer in the mitochondrial genome of P60002, as the majority of this genome was identical to Clade I (P60002 – regions A and C), but a 4-kb segment (ChrM: 19000-22000) encoded 34 polymorphisms that were identical to Clade II and were entirely absent in Clade I (P60002 – region B, Fig. 6B and Fig. S13). We also found that maximum parsimony approaches mostly separated the P90145 mitochondrial genome into Clade II and SA regions and the P60002 mitochondrial genome into Clade I and II regions (Fig. 6D), consistent with visual alignments (Fig. 6A,B). Direct Sanger sequencing of the mitochondrial genome (regions 5000-5700,18500-19500 and 29000-30000) supported the SNP designations from whole genome sequencing and therefore establish that recombination has occurred within the mitochondrial genomes of P60002 and P94015 (Fig. 6E, S14).

**Figure 6.**
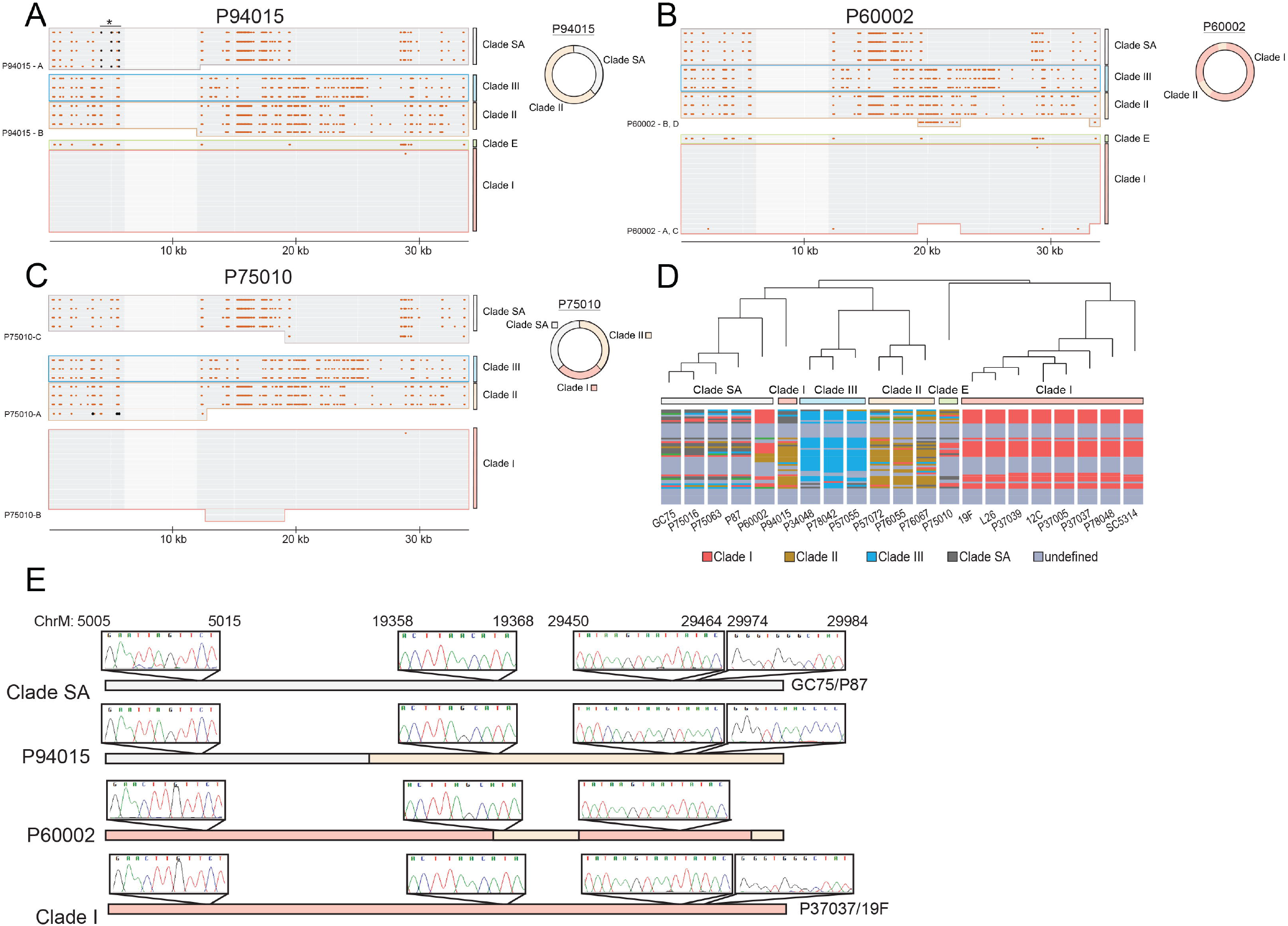
Mitochondrial genomes in *C. albicans* display recombinant genotypes. The mitochondrial (mt) genome sequences of 21 clinical isolates of *C. albicans* were compared. The positions of SNPs that differ from the SC5314 assembly are shown excluding ChrM:35000-41000 due to the absence of any SNPs in this region. The mt genomes for P94015 (**A**), P60002 (**B**), and P75010 (**C**) are highlighted to show the mosaic configuration of SNPs relative to other clades. The mt genome in P94015 contains regions that align with those of Clade II and Clade SA (clade-specific pattern marked with an asterisk with key positions shaded), whereas the P60002 mt genome aligns with sequences for Clades I and II. A 6 kb region devoid of SNPs is more lightly shaded. The three clade SA-specific SNPs in the region that demonstrate alignment of P94015-A are colored black to aid visual alignment. **D.** The similarity of mt regions from each isolate was compared by analyzing 2 kb windows from each strain relative to consensus SNP patterns for each clade. The window was color-coded to designate the clade with greatest similarity. dark grey-SA, blue-Clade III, mustard-Clade II, red-Clade I, light grey-no clade consensus. **E.** Two strains from Clade SA and Clade I along with P60002 and P94015 were Sanger sequenced across three separate 1 kb regions of their mitochondrial genomes. Chromatograms highlighting variant positions consistent with recombination between clades producing the SNP patterns present in P60002 and P94015 are shown along each chromosome.

We further note that the single representative of Clade E in our collection, P75010, also displays strong evidence of recombination in its mitochondrial genome (Fig. 6C). The first ~12 kb of P75010 (P75010-A) aligns closely with Clade II and encodes all three clade II-specific SNPs in this region. The following ~7 kb region of P75010 (P75010-B) matches that of Clade I strains, and is followed by a region with clear homology to Clade SA, encoding 22 of 25 Clade SA-specific SNPs. Identity to several clades infers that multiple recombination events have given rise to this complex SNP pattern. Taken together, these results reveal recombination events have occurred within *C. albicans* mitochondrial genomes and provide clear evidence that sexual/parasexual processes have occurred during *C. albicans* evolution.

## Discussion

Our analysis of *C. albicans* genome structure reveals a number of important aspects concerning mutational patterns during natural evolution of the species. We highlight that (1) non-coding and heterozygous regions of the genome accumulate more mutations than coding and homozygous regions, respectively, (2) there is a significant association between the positions of emergent SNPs and indels, (3) diverse LOH events contribute to genetic inheritance, including long-tract LOH events that extend to the ends of the chromosomes, (4) there is evidence for selection acting on natural populations, (5) a subset of strains exhibit mosaic nuclear and mitochondrial genomes, and (6) analysis of specific chromosomal regions reveals clear evidence for inter-clade recombination.

### Mutations driving natural evolution of the *C. albicans* genome

Mutation rates vary across eukaryotic genomes in a context-dependent manner [44, 45]. We found that mutation patterns arising in natural *C. albicans* populations similarly exhibit a non-random distribution across the genome. Our analysis focused on the distribution of strain-specific SNPs, as these SNPs are likely to have emerged since these strains diverged from one another. We found that the location of strain-specific SNPs was biased towards heterozygous over homozygous regions of the genome. This is consistent with recent studies that showed that mutations arising during meiosis occurred more frequently within heterozygous than homozygous regions of eukaryotic genomes, although the mechanism driving this bias is unknown [46, 47]. Our results extend these findings by indicating preferential accumulation of mutations within heterozygous regions during mitotic growth in *C. albicans*, suggesting that elevated mutation rates may be a common feature associated with heterozygosity.

Our analysis also reveals that emergent SNPs and indels cluster together within the *C. albicans* genome, with a significant enrichment of SNPs within 10 bp of emergent indels. This is similar to what has been observed in other eukaryotic species where indels were found to promote elevated substitution rates close to the founding indel [34-36, 48, 49]. Moreover, the transition:transversion ratio was significantly lower at these mutation sites compared to the genome average. This is consistent with a model in which the recruitment of error-prone polymerases results in increased mutation rates during indel formation [35].

Given that recombination events can be mutagenic [37, 40, 50], we also examined whether *de novo* SNPs were more prevalent close to the breakpoints between heterozygous/homozygous regions. However, we found that the regions flanking the borders of *C. albicans* LOH tracts encoded fewer mutations than the genome average. Other studies have similarly found varying associations between recombination crossovers and mutation rates; some recombination break points exhibited no effect on mutation rates [51], some showed increased rates [47], and some displayed reduced rates [52], similar to the current analysis in *C. albicans*.

The location of strain-specific polymorphisms was biased towards non-coding segments of the *C. albicans* genome. Overall, 51.2% of strain-specific SNPs and 85.0% of strain-specific indels resided in non-coding regions, despite these regions representing just 36.7% of the genome. This indicates that selection is likely limiting both SNP and indel mutations from accumulating in protein-coding sequences in *C. albicans*. Where mutations did occur within genes, these were biased towards certain gene classes and mutation types. Genes containing the highest SNP frequencies encoded snoRNAs, consistent with studies in other eukaryotes [53]. High expression of these noncoding RNAs makes them particularly vulnerable to mutations during collisions between DNA and RNA polymerases [54] and these genes are also generally more accepting of mutations than protein-coding genes [55].

### Evolutionary impact of loss of heterozygosity

Loss of heterozygosity events are a frequent occurrence in *C. albicans* genomes [1, 56]. Previous analysis of the 21 *C. albicans* genomes noted that LOH tracts can be highly variable in size, with several isolates containing chromosomes that had experienced whole chromosome LOH [28]. Here, we made the observation that the vast majority of very large LOH tracts (defined as LOH tracts >50 kb that were not whole chromosome LOH events) initiated between the centromere and the telomere and extended all the way to the chromosome ends. Such LOH events are likely due to break-induced replication, in which one chromosome homolog is used as a template to repair a double-strand DNA break in the other homolog, although reciprocal crossover events can also generate long LOH tracts [26, 57, 58]. The breakpoint for most of these long-tract LOH events varied between individual strains suggesting that independent LOH events had occurred. Short-tract LOH (<50 kb) was also common with approximately of half of these events being shared between related strains (Fig. 3, Fig. S8, Table S5), indicating that homozygosis of certain regions may confer a selective advantage or are not sufficiently deleterious to be selected against. LOH of specific alleles has been shown to alter phenotypes that impact a range of *C. albicans* traits from growth rates to virulence to drug resistance [29, 59, 60].

The cumulative effects of all of these mutational forces on *C. albicans* genomes are accelerated evolution of heterozygous regions relative to homozygous regions. Eventually, the rapid accumulation of deleterious mutations during clonal growth would be expected to result in a fitness decline due to Muller’s ratchet [61, 62]. The occurrence of LOH may counterbalance these forces both by culling mutations from the genome and by reducing heterozygosity to slow evolutionary rates across the genome. LOH events appear at different points during the evolution of individual strains as most LOH tracts are not shared even between closely-related isolates. Indeed, accumulation of emergent strain-specific SNPs within more ancestral LOH tracts demonstrates that these LOH events are not recent occurrences and have further mutated since their origin.

### Evidence for genetic exchange in clinical *C. albicans* isolates

Studies of *C. albicans* population structure point to a largely clonal mode of reproduction, yet there is also evidence of mixed evolutionary histories indicative of a sexually/parasexually reproducing species [21, 22, 24, 27, 63-65]. Furthermore, genes encoded at the mating locus show evidence for ongoing selection consistent with a conserved role in regulating sexual/parasexual reproduction [66]. In this study, we interrogated whole genome data for evidence of genetic admixture, while noting that LOH events can complicate analysis of recombination in diploid strains (Fig. 4, S10). We reveal that a subset of isolates contain mosaic genomes, consistent with these genomes being the products of mating between different *C. albicans* clades. Both nuclear and mitochondrial genomes of P60002 and P94015 show recombinant genotypes supporting a (para)sexual origin for these strains. Genetic information from multiple clades contributed to these genomes and recombination tracts varied in length from a few kb to hundreds of kb (Fig. 5 and 6). The existence of a subset of *C. albicans* strains with mosaic genomes is similar to what has been observed in wild and domesticated strains of *S. cerevisiae*, where both non-mosaic and recombinant mosaic genomes have been identified [67]. Analysis of admixed genomes in P60002 and P94015 suggests that recombination events may be relatively ancient, as recombination involves multiple clades and more recent mutational events have obscured the precise evolutionary histories of these strains.

Disagreement between strain phylogeny by MLST and Ca3 fingerprinting may also be indicative of recombination in the population. Based on Ca3 fingerprinting, P94015 should cluster with other Clade I strains and is supported by MLST analysis, which groups P94015 and other MLST 6 strains closer to MLST 1 / Clade I than other groups [28]. Yet, whole genome sequencing clusters P94015 closest to Clade SA (and even Clade II and Clade III strains) than other Clade I strains. This could reflect how recombination can distort the phylogenetic relationship between strains when based on analysis of a small subset of loci. Analysis of additional *C. albicans* isolates will help define how prevalent recombination is in the species and whether these recombination events are ancient or more recent occurrences.

Critically, we were able to identify regions in *C. albicans* genomes that exactly matched the pattern of recombinant SNPs expected from mating events between two extant clades. This was exemplified by one region in P94015 which consisted of multiple SNPs that exactly matched those present in Clade I followed, after a short gap, by a run of SNPs that precisely matched those in Clade SA (Fig. 5B). Recombination events were also clearly evident in mitochondrial genomes of at least 3 of the 21 isolates examined (P60002, P75010 and P94015). Taken together, these studies provide the clearest evidence to date that *C. albicans* populations have been shaped by (para)sexual exchange.

In summary, *C. albicans* genomes reveal multiple signatures of the forces that have shaped genetic diversity within the species. Both short and long LOH events have played a major role in increasing population diversity, with large tracts extending along the terminal regions of many chromosome arms that impact hundreds to thousands of polymorphisms. Base-substitution mutations and indels cluster within heterozygous regions of the genome, suggestive of faster evolution of these regions, and recombination between isolates has generated mosaic nuclear and mitochondrial genomes with potentially profound consequences for adaptation. The diploid heterozygous genome of *C. albicans* is therefore a highly dynamic platform on which selection can act.

## Materials and Methods

### Variant calling, Processing, and Display

Whole genome sequencing, variant identification, and Loss of Heterozygosity (LOH) windows were identified in a previous study (Hirakawa et al. 2015). Briefly, BWA 0.5.9 [68] read alignments were filtered with a minimum mapping quality of 30 using SAMtools [69]. To reduce the incidence of false positive SNPs called near indels, poorly aligned regions were realigned using GATK RealignerTargetCreator and IndelRealigner (GATK version 1.4-14) [70]. Both prior SNP variants and variants present in the mitochondrial genomes and indels for both mitochondrial and nuclear genomes described here used GATK UnifiedGenotyper, and filtered using the GATK VariantFiltration using hard filters (QD<2.0, MQ<40.0, FS>60.0, MQRankSum<-12.5, ReadPosRankSum<-8.0). The genome sequences used in this study are available under BioProject ID PRJNA193498 (http://www.ncbi.nih.gov/bioproject). SNP data is available from dbSNP (http://www/ncbi.nih.gov/projects/SNP) under noninclusive ss# 1456786277 to 1457237021. SNPs were called homozygous if greater than 90% of the reads contained the non-reference nucleotide. This high threshold also reduced miscalling due to trisomic chromosomes/chromosomal regions.

To assess heterozygous and homozygous variant calls, the number of reads at each variant position was divided by the total number of reads at that position in that strain. On average, each variant position had 52.17 reads with an interquartile range from 33 to 71 reads. The distribution of the allelic ratio at each variant is plotted in Figure S15. The mean number of reads of the allele divided by the total number of reads for heterozygous positions was 0.4499 and for homozygous regions, 0.9889. Thus, a 90% threshold was used for homozygous regions whereas heterozygous regions span from 10-50%.

From this dataset, strain-specific SNPs and indels were parsed into a separate set for additional analysis. Strain-specific variant features were required to be uniquely called in only one of the 21 strains at its genomic position. The data sets are available online in a searchable interface, using R shiny for the backend: [snpmap.asc.ohio-stagte.edu]. Manual interrogation of 100 variants using the Integrated Genome Viewer [71] confirmed the variant presence and call quality metrics in all 100. Manual interrogation of SNP-indel pairs in the pileup confirmed 98% (49 of 50) are valid using the same criteria.

To adjust for mutations in homozygous regions that may have occurred but then lost due to LOH, the following calculation was used for each homozygous region: ((homozygous SNPs/all SNPs)+1)*homozygous SNPs)/length. This effectively doubles the number of homozygous SNPs in hom regions to account for heterozygous SNPs that are lost due to LOH. Even with this doubling, significantly more emergent SNPs occurred in het regions than in hom regions.

### Phylogenetic construction of strain relatedness

The methods used to construct the phylogeny of the sequenced strains was previously described in Hirakawa *et al*. [28]. Briefly, the phylogenetic relationship of strain relatedness was constructed using all whole genome SNP calls, which totaled 113,339. A distance based tree was estimated relying on maximum parsimony and a stepwise matrix where homozygous positions are two steps away compared to one step for heterozygous positions. SNP positions were resampled in 1,000 bootstrapped samples and each node indicates the bootstrap support.

### Determination of LOH

Previously defined heterozygous and homozygous genomic region were used as detailed in [28]. Briefly, the SNP density of homozygous and heterozygous positions was calculated across the genome in non-overlapping 5 kb windows for each isolate. The resulting 5 kb windows were managed as follows:1) Homozygous regions shared between individual strains and SC5314 were identified and marked as homozygous; 2) a single 5 kb window adjacent to these regions lacking any polymorphisms was merged into homozygous tracts if present; 3) contiguous, adjacent windows with a significantly higher frequency of homozygous SNPs than SC5314 homozygous regions (>0.4 SNPs/kb) were merged allowing one intervening window lacking sufficient polymorphism into homozygous tracts. These regions were defined as homozygous whereas remaining regions covered by 5 kb windows of the genome were designated heterozygous and contained significantly more heterozygous SNPs. The borders between homozygous and heterozygous regions were manually inspected for accuracy.

### Introgression/Tree Violations

Two independent methods were used to assign clade designations for nuclear and mitochondrial genomes in non-overlapping 25,000 or 750 bp windows, respectively. The first process assigned the clade that best resembled the query strain’s SNP pattern. However, Clade I contains fewer SNP calls due to alignment to the SC5314 reference genome (also a Clade I isolate) that can introduce an artificial bias. Therefore, a dataframe was constructed where each row represented any SNP contained within any strain in the query window. A SNP could only be counted as a single row so the identity of that SNP position was recorded as “0” for the absence of the SNP, “1” for a heterozygous position, and “2” for a homozygous position. The correlation between the target strain’s resulting numeric “SNP” profile and each other strains was individually calculated. Scores from strains within the same clade were averaged and taken as the absolute value: *Clade score* = *abs*(*mean*(*cor*(*query strain SNP, strain X SNP*))). The clade with the highest similarity score (and thus greatest proportion of shared SNPs) was selected as the most similar clade for that window. This process was repeated across the full genome. As a followup, this process was repeated with removing all strains within the query strain’s clade and the scores recalculated.

The second phylogenetic approach used an expression matrix listing the 21 strains against all possible SNP positions present for each non-overlapping window. For each cell, we denoted a 0 if the respective SNP was not present in the respective strain, a 1 if one copy of the SNP was present, and a 2 if both copies were present. A distance matrix using a binary method (R dist) was constructed from this data and finally resulted in a phylogenetic tree (R hclust). The appropriate number of K clusters for the phylogenetic output of each window was estimated by traversing through all 21 possible values (1 to 21) incrementally using R cutree. The first k-value was chosen that allowed for the target strain to cluster with at least two other strains that were members of the same clade as defined by the current phylogenetic relationship among the sequenced strains. These criteria effectively eliminate Clade E because it only contains one strain. Additionally, this approach was assessed for congruence manually across candidate regions.

### Sanger sequencing

The association between SNPs and indels identified in the NGS data was tested by amplification of specific regions that were either shared among a number of strains (P75063) or strain-specific (P60002). Two regions were PCR amplified using primers AGTCGGTGATGTCTATAGTG / GCTGTCCTTGGATCATTGAT to amplify Chr7:48017..48664 in P75063 and TTCTGCTGTTGCTGCTGCTA / CTGTCAACTGTCAACCAAAG to amplify ChrR: 19979596..1998145 in P60002. Amplicons were purified and Sanger sequenced.

Mitochondrial SNPs were verified using 3 sets of primers to amplify different regions of ChrM across the 21 natural isolates. Two isolates were analyzed from each clade including P60002, P75010, and P94015 strains. Primers TTAGTAGTGTCGGTGTCTTC / AGAGAGGGTTTTGGTTAGGG were used to PCR amplify ChrM: 4899..6076, GAATCTCAGAGACTACACGT / GTGGTATACGACGAGGCATT were used for ChrM: 18265..20660, and TGGGAAGTAGAGGCTGAAGA / AGGGGCATTATAAGGAGGAG were used for ChrM: 28094..29548. PCR amplicons were purified and Sanger sequenced.

### Statistical Testing

Statistics were performed as Student’s T-test unless otherwise indicated. All statistical tests were performed in the R 3.2.5 programming environment [72].

## ACKNOWLEDGEMENTS

We thank Iuliana Ene and Christophe D’Enfert for comments on the manuscript, Christina Cuomo for technical advice, and members of the Bennett and Anderson labs for helpful discussions. This work was supported by National Institutes of Health grants AI081704 and AI122011 (to R.J.B.), a PATH award from the Burroughs Wellcome Fund (to R.J.B.), and by a Karen T. Romer Undergraduate Teaching and Research Award (to J.M.W.).

## Supporting Information Legends

**Figure S1. The distribution of polymorphisms is non-random.** The number of strains encoding each SNP (A) or indel (B) was determined and the frequency plotted. A best-fit line (red) was plotted for each distribution.

**Figure S2. LOH can obfuscate inheritance by descent patterns of shared polymorphisms.** LOH of opposing alleles in a common ancestor can make it appear that mutations arose independently despite mutations sharing a common origin. LOH of homolog A in Clade III produces a different SNP pattern than that of Clade II although both arose from the same ancestral strain.

**Figure S3. Number of polymorphisms correlates with branch length.** The branch length on the phylogenetic tree to the nearest node was correlated against the number strain-specific SNPs (**A**) or strain-specific indels (**B**).

**Figure S4. Transitions are present more frequently than transversions during strain evolution.** The percentage of SNPs that result in transitions and transversions was calculated both for all SNPs and for strain-specific SNPs.

**Figure S5. Characterization of indels across *C. albicans* isolates.** The number of strain-specific indels was plotted for either intergenic (**A**) or genic (**B**) mutations based on the indel size, ranging from 1-10 nucleotides. Blue indicates deletions and yellow indicates insertions.

**Figure S6. Verification of SNP-indel association by Sanger sequencing.** Four regions that contained SNPs tightly linked to indels were chosen to be assessed by Sanger sequencing. The genomic DNA from Chr7 in strain P76055 (**A**), Chr5 in GC75 (**B**), Chr7 in P87 (**C**), and Chr2 in P57072 were Sanger sequenced and encoded the putative SNPs and indels (colored boxes) as expected from whole genome sequencing. The reference SC5314 sequence is shown for comparison to the Sanger sequenced DNA below along with the chromatogram aligned to the polymorphisms identified from whole genome sequencing. Below each schematic are listed the informative sites.

**Figure S7. Associations of strain-specific variants corrected for clade bias.** Strains-specific mutations from 12 strains (Clade I: 12C, L26, P78048; Clade II: P57072, P76055, P76067; Clade III: P34048, P78042, P57055; Clade SA: P87, GC75, P75063) were retained to ensure that clade representation did not bias these associations. **A.** The relative frequency of strain-specific SNPs (blue) and indels (orange) was plotted across the genome using 5000 bp sliding windows. Distance to the nearest SNP for each indel was measured in a 100 bp window (with non-overlapping 5 bp intervals) surrounding the indels for either strain specific (**B**) or all (**C**) variants.

**Figure S8. Mapping of heterozygous and homozygous regions of *C. albicans* genomes.** Schematic showing heterozygous (red) and homozygous (blue) regions of sequenced ***C. albicans*** isolates. Note that most long-tract LOH events (>50 kb) start between the centromere and the telomere and extend to the ends of the chromosomes. Chromosomes are displayed along the bottom with green circles indicating centromeres, blue boxes denoting major repeat sequence (MRS) loci, and the orange box signifying the *MTL* locus.

**Figure S9. *C. albicans* SNP patterns generally follow inheritance by descent.** Polymorphisms for the set of *C. albicans* genomes are plotted for two loci that display SNP patterns consistence with inheritance by descent. Heterozygous and homozygous SNPs are purple and orange, respectively.

**Figure S10. LOH alters inheritance patterns of *C. albicans* polymorphisms.** Polymorphism patterns are plotted for two loci (**A, C**) that do not follow inheritance by descent. Similar genotypes are color-coded and connected to each other. Cartoons depicting LOH of heterozygotes in opposing directions to produce the observed SNP pattern (**B, D**) provide the most parsimonious explanation for those two loci. Heterozygous and homozygous SNPs are purple and orange, respectively.

**Figure S11. Modes of inheritance in *C. albicans*.** Both LOH and mating can impact SNP patterns in *C. albicans* strains. Analysis of the distribution of heterozygous and homozygous SNPs can help differentiate between these two possibilities’. Following LOH, all affected SNPs are homozygosed and derived strains will therefore contain homozygous variants of homolog A or homolog B. LOH can therefore make the precursor to these LOH events (Strain B in the figure) appear ‘recombinant’, as it will contain heterozygous SNPs at positions that are homozygous in the lineages that underwent LOH. In contrast, strains that are related due to recombination may share only heterozygous SNPs between lineages or may share a mix of heterozygous and homozygous SNPs, with homozygous positions due to inheritance of the same SNP from both parents (see also Figure 3B). These patterns may be further complicated by additional short-tract LOH events, as indicated on the right of the figure.

**Figure S12. Evidence for recombination in two *C. albicans* strains.** The SNP patterns for two regions highlight recombinant genotypes in strains P60002 (A) and P94015 (B). The DNA segments corresponding to different clades are aligned next to the appropriate group and labeled according to homologous tracts. Heterozygous and homozygous SNPs are purple and orange, respectively. **C.** Consensus SNP patterns for each clade were used to assess genome similarities between all isolates in 50 kb sliding windows. In this case, all strains from the same clade were removed to force the closest match for each window to be assigned by color-coding to the most similar clade. The SNP patterns for two strains, P60002 and P94015, stand out with respect to their position in the constructed phylogenetic tree and assigned clade. dark grey-SA, blue-Clade III, mustard-Clade II, red-Clade I, light grey-no clade consensus.

**Figure S13. Alternative mode of recombination in the P60002 mtDNA genome.** The mitochondrial genome of P60002 may be the result of recombination between clade SA (P60002-B,E), clade I (P60002-A,C,F), and clade II (P60002-D,G), although we note that the resolution of SNP patterns cannot distinguish all clades (e.g., regions B, E have identical SNP patterns in both clade SA and clade I).

**Figure S14. Verification of mitochondrial SNPs by Sanger sequencing.** Two strains from Clades I, II, III and SA, as well as P60002, P75010, and P94015 were sequenced across three separate 1 kb regions of the mitochondrial genomes to assess variant calls from whole genome sequencing. Chromatograms highlighting variant positions consistent with recombination between clades producing the SNP patterns present in P60002 and P94015 are shown along each chromosome.

**Figure S15. Read depth of variant calls define heterozygous and homozygous positions.** The proportion of reads encoding a variant at each position was calculated relative to the total number of mapped reads at that position. Plotting each position for all strains produced a distribution that was separated at 90% of reads encoding a variant allele. The distribution of variant position <90% were defined as heterozygous and those >90% as homozygous.

**Figure S16. Positioning of SNPs and indels across *C. albicans* genomes.** A. The relative frequency of all SNPs (blue) and indels (orange) was plotted across the genome using 5000 bp sliding windows. **B.** Number of SNPs within 100 bp of each strain-specific indel was plotted. The average number of SNPs in an equal number of random 100 bp windows bootstrapped 1000 times is shown (red line). **C.** The distance of the nearest SNP to all indels for each strain pair was plotted. (Compare to association of strain-specific SNPs and indels in Figure 1C).

**Figure S17. Associations of strain-specific variants with genic content when corrected for clade bias.** Strains-specific mutations from 12 strains (Clade I: 12C, L26, P78048; Clade II: P57072, P76055, P76067; Clade III: P34048, P78042, P57055; Clade SA: P87, GC75, P75063) were retained to ensure that clade representation did not bias these associations. The ratio of genic to intergenic strain-specific SNPs (**A**) and indels (**B**) was calculated for each of the 12 isolates and falls below the fraction of the genome that encodes protein-coding genes (red line). **C.** The density of strain-specific SNPs from these 12 isolates was measured for all genes in the *C. albicans* genome and plotted against the average SNP density for thesel isolates (red line). **D.** Genes enriched for strain-specific SNPs were identified and functional enrichment was assessed using GO term analysis. snoRNAs were significantly enriched for strain-specific SNPs in these 12 isolates. **E.** The placement of the SNPs was determined by breaking snoRNAs into five equal segments. SNPs were significantly enriched in the 5’ end of the RNAs.

**Table S1. Percentage of the genome encoding SNPs among the sequenced *C. albicans* isolates.**

**Table S2. Strain-specific SNPs among sequenced *C. albicans* isolates.**

**Table S3. Strain-specific indels among sequenced *C. albicans* isolates.**

**Table S4. Genes with significant enrichment of strain-specific SNPs. Gene lists are provided at three different SNPs/nt cutoffs.**

**Table S5. Heterozygous and homozygous regions for the sequenced *C. albicans* isolates.**

